# A potent antibiofilm agent inhibits and eradicates mono- and multi-species biofilms

**DOI:** 10.1101/2020.03.25.009126

**Authors:** Lexin Long, Ruojun Wang, Ho Yin Chiang, Yong-Xin Li, Feng Chen, Pei-Yuan Qian

**Affiliations:** SZU-HKUST Joint PhD Program in Marine Environmental Science, Shenzhen University, Shenzhen, China; Department of Ocean Science and Hong Kong Branch of Southern Marine Science and Engineering Guangdong Laboratory, Guangzhou, The Hong Kong University of Science and Technology, Clear Water Bay, Kowloon, Hong Kong, China; Department of Chemistry, The University of Hong Kong, Pokfulam, Hong Kong, China; Institute for Advanced Study, Shenzhen University, Shenzhen 518060, China

**Keywords:** Elasnin, Biofilms, Biofouling, Infections, Natural products

## Abstract

Biofilms are surface-attached multicellular communities that create many problems in human health and various industries. Given the prominence of biofilms in biofouling and infectious diseases, antibiofilm control approaches are highly sought after. In the present study, we identified elasnin as a potent antibiofilm agent through a bioassay-guided approach. Elasnin specifically inhibited the biofilm formation of bacterial mono-species and eradicated the mature biofilm of Gram-positive bacteria at concentrations below 2.5 μg/mL with a low toxic effect on cells and a low resistance risk. Confocal observations illustrated that elasnin decreased cell aggregations and destroyed the biofilm matrix. Furthermore, elasnin-based antibiofilm coatings were prepared and inhibited the formation of multi-species biofilms and the attachment of large biofouling organisms in the field test. These findings suggest that elasnin is a promising antibiofilm agent for future applications in biofilm control.

**Importance:** Due to the increased diversity of biofilm-associated infections and the failure of conventional antimicrobial treatment, new and effective biofilm-specific pharmacologic strategies are urgently needed. Elasnin is a new antibiofilm natural product produced by Streptomyces with high efficiency and low toxicity. Elasnin effectively destroyed the biofilm matrix of Gram-positive bacteria, thus making them more susceptible to antibiotics. Unlike currently deployed antibiotic vancomycin, which exclusively targets essential life processes and kills the pathogen, elasnin did not exhibit bactericidal effect and thus held great potential in delaying resistance. With high yield, elasnin-based coatings were easily prepared with low expenditures and exhibited favorable performance in field test. Collectively, the antibiofilm properties of elasnin, combined with the low cost of supply and the low risk of resistance, could provide the basis for the development of a novel antibiofilm agent that could help fight to antibiotics resistance.

## Introduction

A biofilm is a microorganism community attached to a surface^1^. It consists of microbial cells massed in the matrix of extracellular polymeric substances (EPS), which contain a large variety of biopolymers such as proteins, nucleic acids, lipids, and other substances^2^. Biofilms can be made up of a single microbial species or multiple species that colonize biotic or abiotic surfaces^3,4^. The elaborate biofilm architecture provides a shield to the microbes in biofilms and offers them the spatial proximity and internal homeostasis needed for growth and differentiation^3–5^. This makes microbial cells much more resistant than their planktonic counterparts to diverse external insults such as antimicrobial treatment, poisons, protozoans, and host immunity^6,7^. For example, biofilms can render organisms 10- to 1000-fold less susceptible to antimicrobial agents; furthermore, organisms in multi-species biofilms are less susceptible to antimicrobial treatment than those in mono-species biofilms due to the more complex interactions^6,8,9^.

Biofilm formation creates many problems in human health and various industries. It is estimated that about 80% of microbial infections were associated with biofilm formation which plays a key role in the pathogenesis and confers on the associated bacteria great resistance to conventional antimicrobial agents^10,11^. The formation of marine biofilms also leads to serious economic loss for various industries such as heat exchange, oil and gas processing, and the maritime industry^12,13^. Physical removal, high-intensity and sustained antimicrobial chemotherapy, and surface coatings are the few antibiofilm strategies available^14^. However, in most cases, these strategies are either costly or toxic to the marine environment. Also, the required high dosage of bactericides for biofilm eradication promotes the development of widespread antimicrobial resistance^12,14,15^. Therefore, efficient, safe, environmentally friendly and cost-effective biofilm control approaches are urgently needed.

*Actinobacteria* are one of the largest taxonomic units in the Bacteria domain and have always been a promising source of antimicrobial compounds. Among the more than 22,000 bioactive compounds discovered by the early 2000s, 45% were produced by *Actinobacteria*^16^. In the present study, we identified an antibiofilm compound elasnin that has low toxicity and potent antibiofilm activities against mono- and multi-species biofilms from an actinobacterial species *Streptomyces mobaraensis* DSM 40847.

## Results

### Bioassay-guided isolation of antibiofilm compounds

Secondary metabolites produced by 12 actinobacterial strains under different culture conditions were first assessed for bioactivity against Gram-positive bacteria (methicillin-resistant *Staphylococcus aureus*) and Gram-negative bacteria (*Escherichia coli*), followed by bioassay-guided fractionation and isolation which led to the identification of four known antimicrobial compounds (e.g., elasnin, xanthone, hitachimycin, and resistomycin) against Gram-positive bacteria. Among these compounds, only the one produced by the strain *Streptomyces mobaraensis* DSM 40847 exhibited antibiofilm activities. This compound was subsequently purified using high-performance liquid chromatography (HPLC) and structurally characterized as a known compound elasnin using ultra-performance liquid chromatography-tandem mass spectrometer (UPLC–MS/MS) analysis and nuclear magnetic resonance (NMR) spectroscopy analysis (Fig. 1 and Fig. S1).

**Fig. 1.**
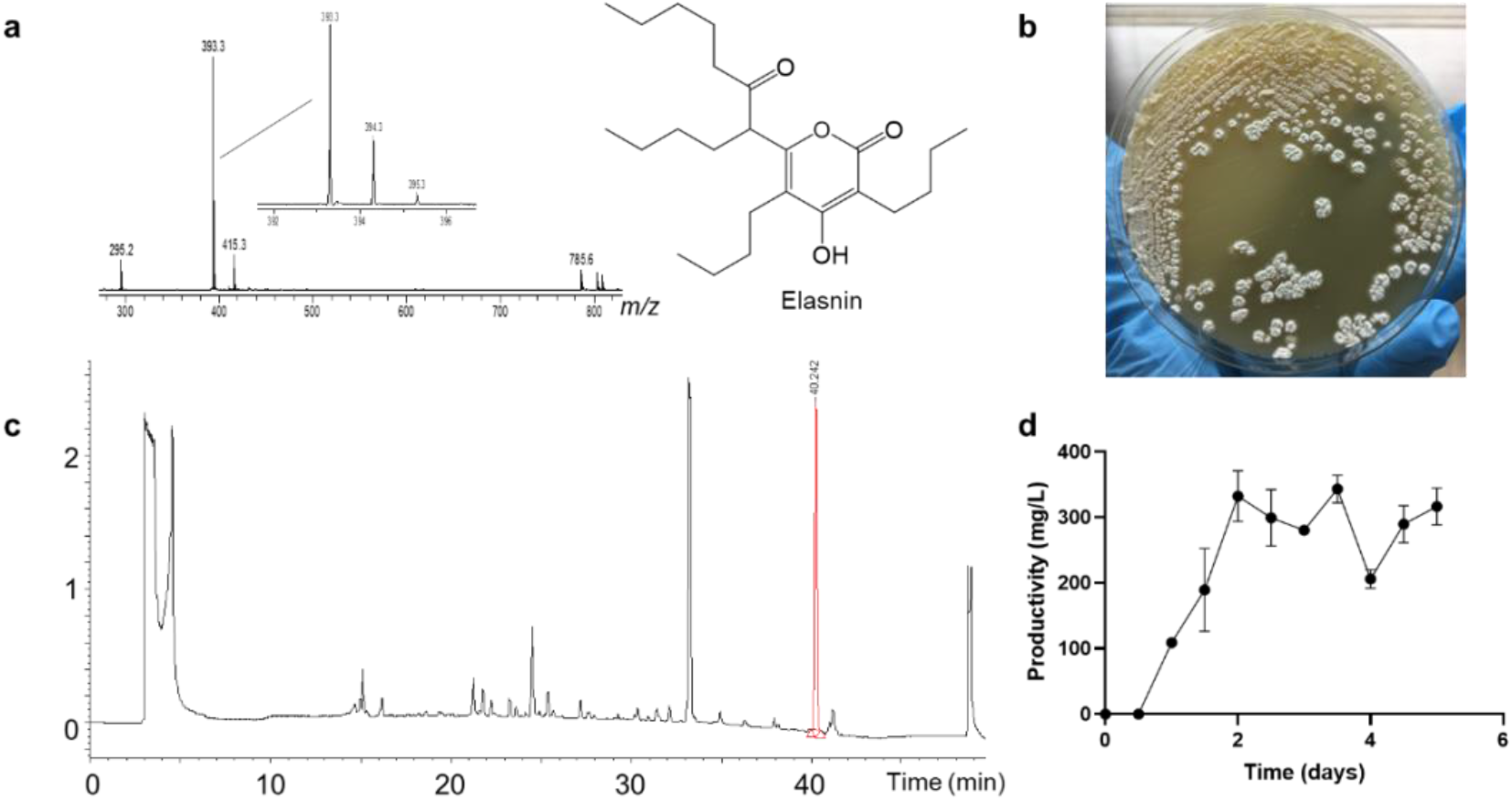
Elasnin was discovered to be produced by *Streptomyces mobaraensis* DSM 40847 with high yield. **a** Mass spectra (ESI) and structure of elasnin. **b** Growth of *S. mobaraensis* DSM 40847 on the GYM (Table S1) agar plate. **c** HPLC analysis of the crude extracts of *S. mobaraensis* DSM 40847. **d** Time course of elasnin’s production in AM4 medium under 30°C.

### Elasnin effectively inhibited biofilm formation and eradicated mature biofilms of MRSA with nonlethal effects on cells and no resistance development

To assess the potency of elasnin against methicillin-resistant *S. aureus* (MRSA), the assays of minimal biofilm inhibitory concentrations (MBICs) and minimal biofilm eradication concentrations (MBECs) were used in this study. Both compounds tested in the present study showed strong biofilm inhibiting activities with MBIC value of 1.25-2.5 μg/mL (Fig. 2a). When bacteria in mature biofilms were challenged with vancomycin and elasnin, vancomycin showed reduced and unstable activity in eradicating biofilms with MBEC (50%) of 10-20 μg/mL (Fig. 2b) while elasnin showed strong eradication activity against mature biofilms with an MBEC (50%) value below 1.25 μg/mL. These results suggest that elasnin’s activities are not affected by the biofilm formation as the effectiveness of vancomycin is obviously influenced.

**Fig. 2.**
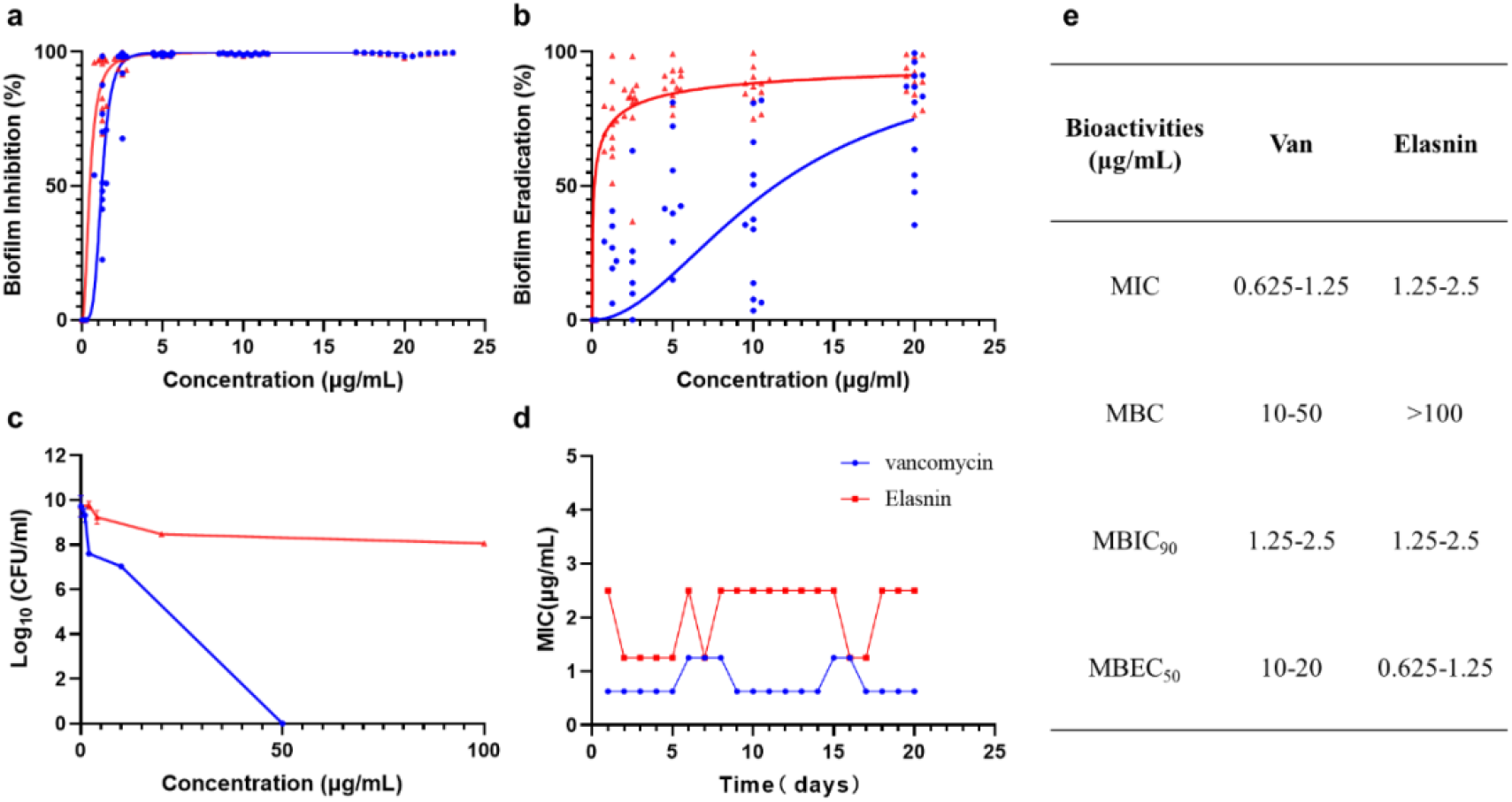
Antimicrobial activities of vancomycin and elasnin against methicillin-resistant *S. aureus* ATCC 43300. **a** Minimum concentration needed for inhibiting 90% biofilm formation. **b** Minimum concentration needed for eradicating 50% mature biofilms. **c** Cell viability after 24 hr treatment of various antimicrobials. **e** resistance development during 20-days serial passaging in the presence of subinhibitory concentrations of antimicrobials. **d** Summary of MICs, MBCs, MBICs, and MBECs. Points below 0% are not shown in the figure.

Planktonic cells of MRSA were susceptible to both two antimicrobial compounds with minimum inhibiting concentrations (MICs) of 0.625-1.25 μg/mL for vancomycin and 1.25-2.5 μg/mL for elasnin. Elasnin exhibited bacteriostatic activity with a minimum bactericidal concentrations (MBCs) exceeding 100 μg/mL, while vancomycin showed a strong bactericidal activity with an MBC between 20 and 50 μg/mL (Fig. 2e). Vancomycin’s bactericidal effect was concentration-dependent, and the MRSA cell density decreased with an increase in compound concentration. In contrast, elasnin showed no lethal effect and the cell viability of MRSA did not significantly differ with the change in compound concentrations (Fig. 2c). Meanwhile, to explore the mode of action of elasnin, we initially attempted to identify resistant mutants of MRSA. Interestingly, MRSA did not develop resistance during continuous serial passaging in the presence of subinhibitory concentrations of elasnin over a 20-day period (Fig. 2d), suggesting a low risk of resistance to elasnin.

### Elasnin destroyed biofilm matrix and released the cells from biofilm

In the present study, we used confocal laser scanning microscopy (CLSM) observations to examine the effect of elasnin on biofilm structures. In biofilm inhibition assay, the untreated biofilms (Fig. 3a) showed distinct shapes with a high density of organized cells and matrix whereas the biofilms treated with elasnin (Fig. 3b) exhibited a large decrease in density of cells and matrix and both of them were randomly distributed. In biofilm eradication assay, pre-formed biofilms were dispersed after treated with elasnin and biofilm cells were released into the media (Fig. 3d). Confocal images demonstrated that the distribution patterns of cells clearly changed after the treatment of elasnin; the untreated biofilm cells (Fig. 3c) distributed as a clump with a rough edge while elasnin treated biofilm cells distributed as narrow strips with smooth edge (Fig. 3d). Similarly, the high density of organized biofilm matrix turned into sparse and scattered after the treatment of elasnin. According to the quantitive analysis, both cells and matrix were significantly reduced after the treatment of elasnin. Compared to the untreated biofilms, the ones treated with elasnin exhibited an around 80% and 35% decrease in density of cells and matrix for biofilm inhibition assay (Fig. 3e); for biofilm eradication assay, the reduction in cells and matrix were over 50% and 70% (Fig. 3f).

**Fig. 3.**
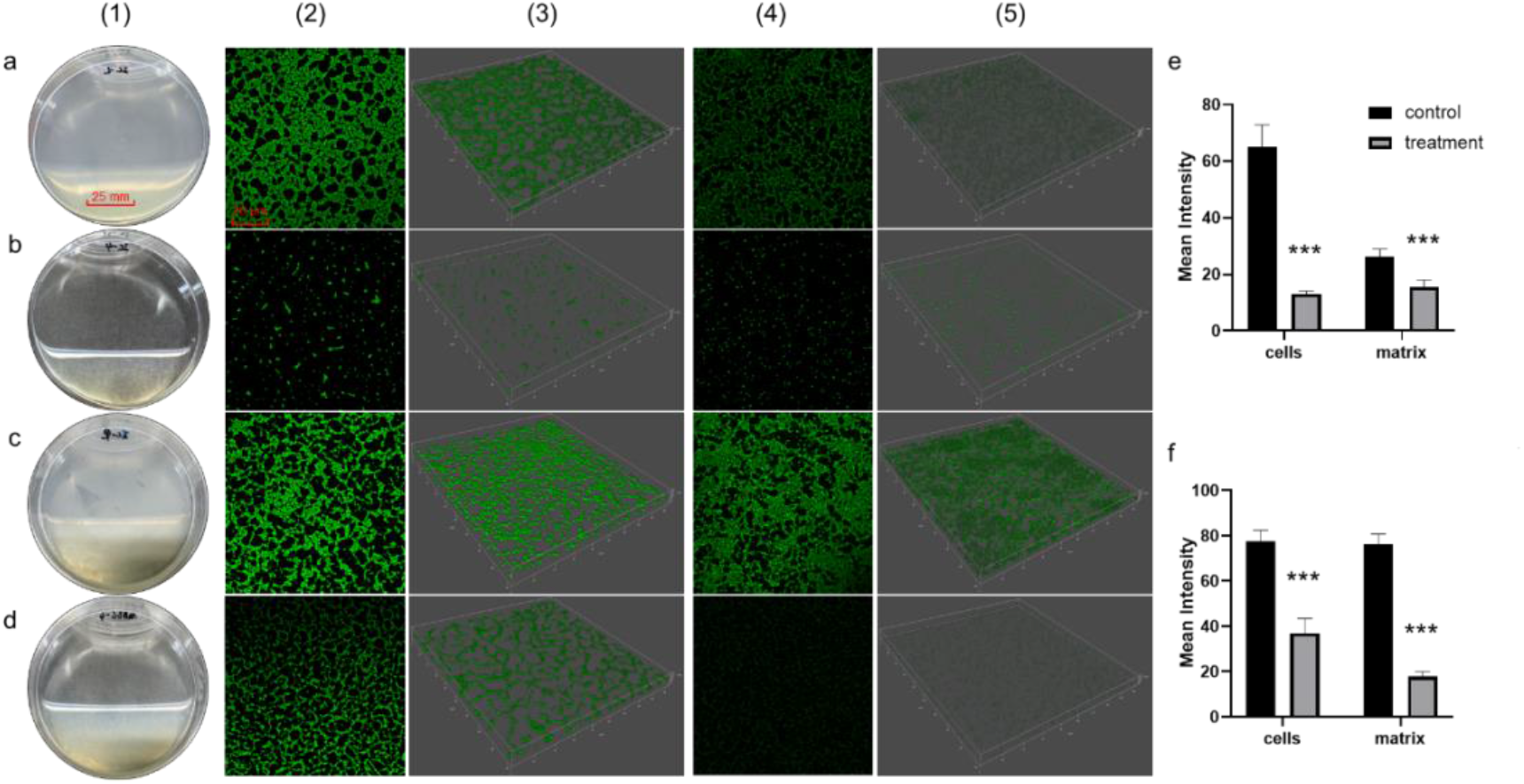
Comparisons between elasnin’s effect on MRSA biofilm cells and matrix. **a** Image of the biofilms after incubation for 24 hr (control). **b** Image of the biofilms after incubation for 24 hr with elasnin treatment at a concentration of 4 μg/mL(treatment). **c** Image of the pre-formed biofilms after another incubation for 18 hr(control). **d** Image of the pre-formed biofilms after another incubation for 18 hr with elasnin treatment at a concentration of 4 μg/mL(treatment). **e** Quantitative analysis of confocal images acquired in biofilm inhibition assay and **f** biofilm eradication assay. Series 1 were pictures of biofilms under direct observation. Series 2 and 3 were two- and three-dimensional confocal images of biofilm cells stained by FilmTracer™ FM® 1-43 green biofilm cell stain. Series 4 and 5 were 2D and 3D images of biofilm matrix stained by FilmTracer™ SYPRO® Ruby Biofilm Matrix Stain. Confocal images were aquired under the same conditions and quantitative analysis was conducted using Leica Application Suite X based on the relative fluorescence of 3D confocal images.

### Elasnin showed preference to the biofilms of Gram-positive bacteria

Among the six strains—*S. aureus* ATCC 25923, *S. aureus* B04, methicillin-resistant *S. aureus* ATCC 43300, *Bacillus subtilis* 168, *E. coli* ATCC 25922 and *Pseudomonas aeruginosa* PAO1—that are used as targets in the antibiofilm efficiency assay, the biofilms of four Gram-positive bacteria were sensitive to elasnin treatment and elasnin’s MBICs ranged from 1.25 to 5 μg/mL (Fig. 4a) while its MBECs ranged from 0.625 to 5.0 μg/mL (Fig. 4b). However, elasnin has no antibiofilm activities against Gram-negative bacteria up to a concentration of 100 μg/mL as shown by the viability of biofilm cells in both the MBIC and MBEC assays, which remained steady regardless of the elasnin concentration used. Therefore, the antibiofilm activities elasnin possess might be specific to Gram-positive bacteria.

**Fig. 4.**
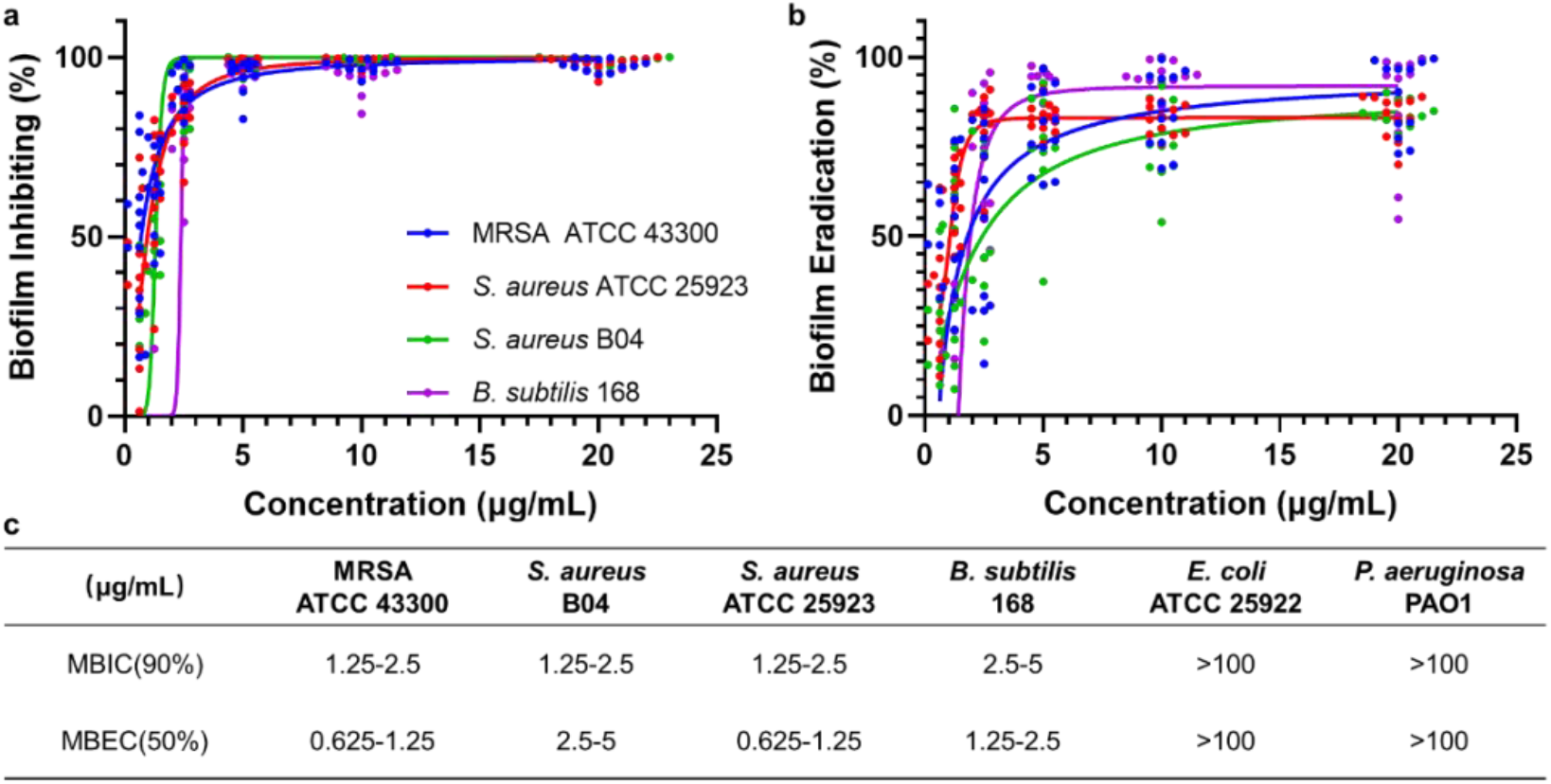
Antibiofilm activity of elasnin against microorganisms. **a** Activities of elasnin to reduce biofilm formation by 90%. **b** Activities of elasnin to eradicate 50% of a mature biofilm. **c** Bacterial strains used in antibiofilm assays and the MBICs/MBECs of elasnin against these strains. Points below 0% and the data of *E. coli* and *P. aeruginosa* are not shown in the figure.

### Elasnin-based antibiofilm coatings inhibited the formation of multi-species biofilms and the attachment of large biofouling organisms in the field test

Given the high antibiofilm efficiency, the low risk of resistance, and the attractive mode of antibiofilm action, elasnin-based antibiofilm coatings (coatings) were prepared and immersed in a fish farm to evaluate this efficiency against natural marine biofilm. Note that in the present study, we used a crude extract of *S. mobaraensis* DSM40847 that contained very high concentrations of elasnin (= 336.64 mg/L in n-hexane, Fig. S2), instead of pure elasnin. The extracts were mixed with polyurethane (polymer) based on poly ε-caprolactone and applied directly on the surface of glass slides. The concentrations of the coatings were calculated based on the percentage of crude extracts in total coatings (polymer and crude extracts) by weight. As such, other compounds in low amounts in the fractionated extract may have exerted an effect on the results of our field testing. However, their effect should be negligible since we did not detect any significant effects of the minor compounds in the crude extracts (Fig. S3).

The release rate of elasnin from the coatings was found to be dependent on both time and concentration for four weeks (Fig. 5c). In general, the release of elasnin from the coatings was keeping at a reasonably low rate throughout the period; the higher the concentration was, the faster elasnin was released into the artificial seawater. The highest release rate of approximately 5 μg day^−1^ cm^−2^ occurred in the second week for the concentration of 10 wt%; for other concentrations, the maximum release rate was around 4 μg day^−1^ cm^−2^ in the first week. The release rate decreased over time and depended on the total amount of elasnin remaining in the coatings. After immersion for four weeks, the release rate dropped to about 1 μg day^−1^ cm^−2^ for the concentrations of 10 wt% and 5 wt% and 0.5 μg day^−1^ cm^−2^ for 1.5 wt% and 2.5 wt%.

**Fig. 5.**
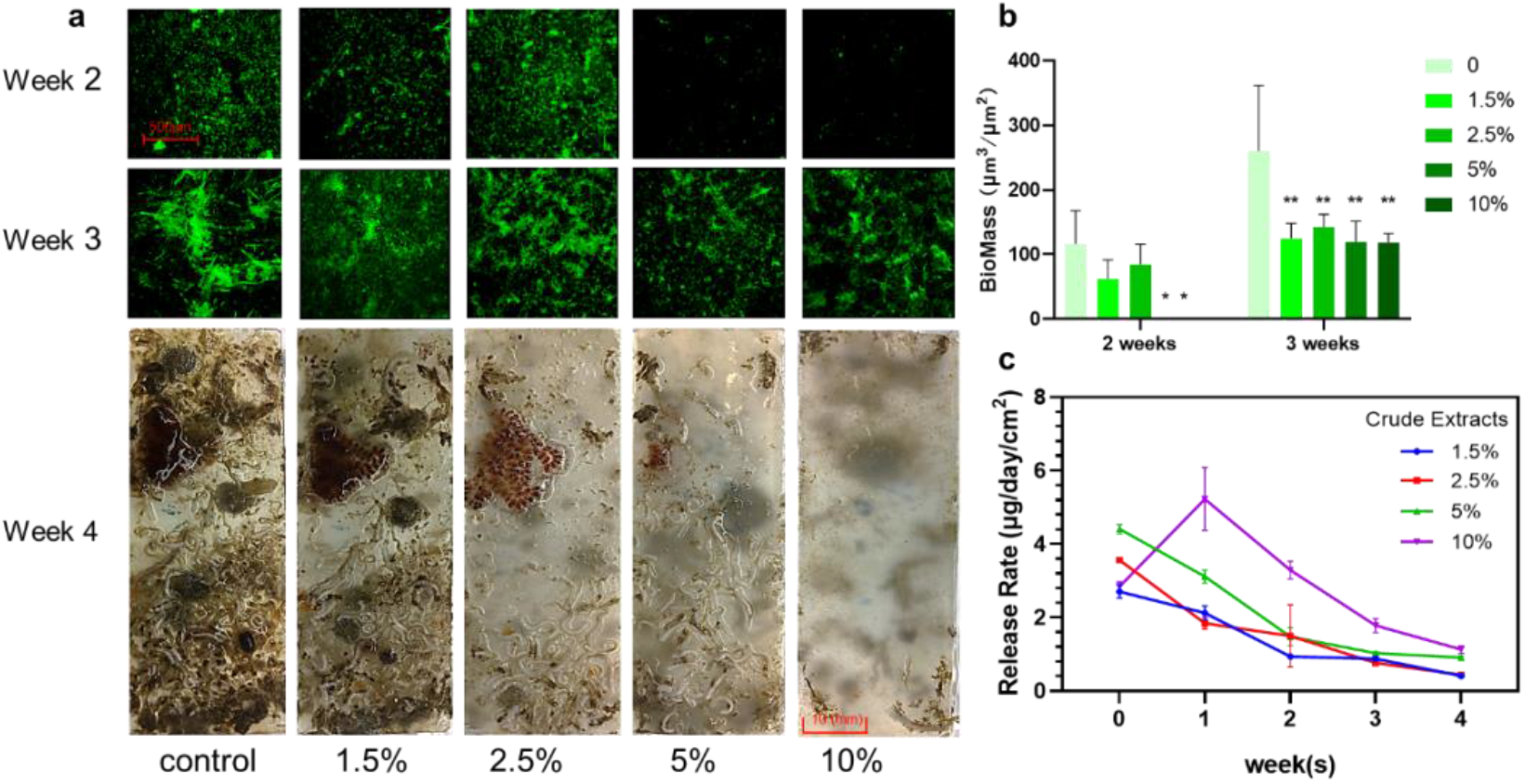
Antibiofilm (weeks 2 & 3) and antifouling (week 4) performance of elasnin-based antibiofilm coatings. **a** CLSM images (weeks 2 & 3) and pictures of the coated surface (week 4). **b** Biomass of biofilms observed by CLSM. **c** Monitoring of elasnin’s release rate in artificial seawater. Biomass was calculated using Comstat 2.1 based on the CLSM images and the values that are significantly different between elasnin-based antibiofilm coatings and control groups are indicated by asterisks as follows: * for p<0.05 and ** for p<0.01.

The performance of the antibiofilm coatings was assayed weekly from the second to the fourth week by direct and CLSM observation. According to the quantitative analysis of CLSM images, the average biofilm biomass on the slides without elasnin was 116.44 μm^3^ μm^−2^ in the second week and 259.95 μm^3^ μm^−2^ in the third week, whereas the average biomass of biofilms measured on 5 wt% and 10 wt% coating slide was less than 0.1 μm^3^ μm^−2^ in the second week and less than 120 μm^3^ μm^−2^ in the third week. For coatings with low concentrations (1.5 wt% and 2.5 wt%), there were no significant differences in terms of average biomass (61.97 μm^3^ μm^−2^ and 84.73 μm^3^ μm^−2^ respectively) in the second week but the biomass was significantly lower than that in the control (259.95 μm^3^ μm^−2^) in the third week, with average biomass of around 125 μm^3^ μm^−2^ and 145 μm^3^ μm^−2^ respectively (Fig. 5b). In the fourth week, slides coated with low concentrations of elasnin (1.5 wt%, 2.5 wt%, and control) were fouled by large marine organisms while the ones coated with high concentrations of elasnin exhibited anti-macrofouling activity and almost no larval settlement, except for a small area near the edges due to edge effects commonly found on testing panels. Overall, elasnin-based antibiofilm coatings inhibited the biofilm formation of multiple bacterial species in the first two weeks. However, after immersion for four weeks, the glass slides coated with low concentrations of elasnin were eventually covered by large biofouling organisms, likely due to the low releasing rate of the elasnin after three weeks.

### Elasnin changed the microbial community structure of natural marine biofilms

Since there was hardly any biofilms developed by the end of the second week but macrofoulers had overgrown by the end of the fourth week, only the three-week-old biofilms developed on 10 wt% coatings and those on the control glass slides (coated with poly ε -caprolactone based polyurethane only) were selected for 16S amplicon analysis to determine the changes in biofilm microbial community triggered by elasnin. A total of 3,000,000 16S rRNA gene sequences (500,000 per sample) were classified into 31 phyla (*Proteobacteria* were classified down to the class level). The microbia composition of the biofilms differed between the 10 wt% coatings and the control slides, as confirmed by alpha- and beta-diversity analysis. In the Bray-Curtis dissimilarity (beta-diversity) dendrogram (Fig. 6a), the control group and treatment group were clustered separately, based on the differences in microbial abundance between samples; the observed OTUs and Shannon diversity for the treated biofilm were significantly lower than those in the control group (Fig. 6b), suggesting that both the species richness and diversity in the treated biofilms were reduced.

**Fig. 6.**
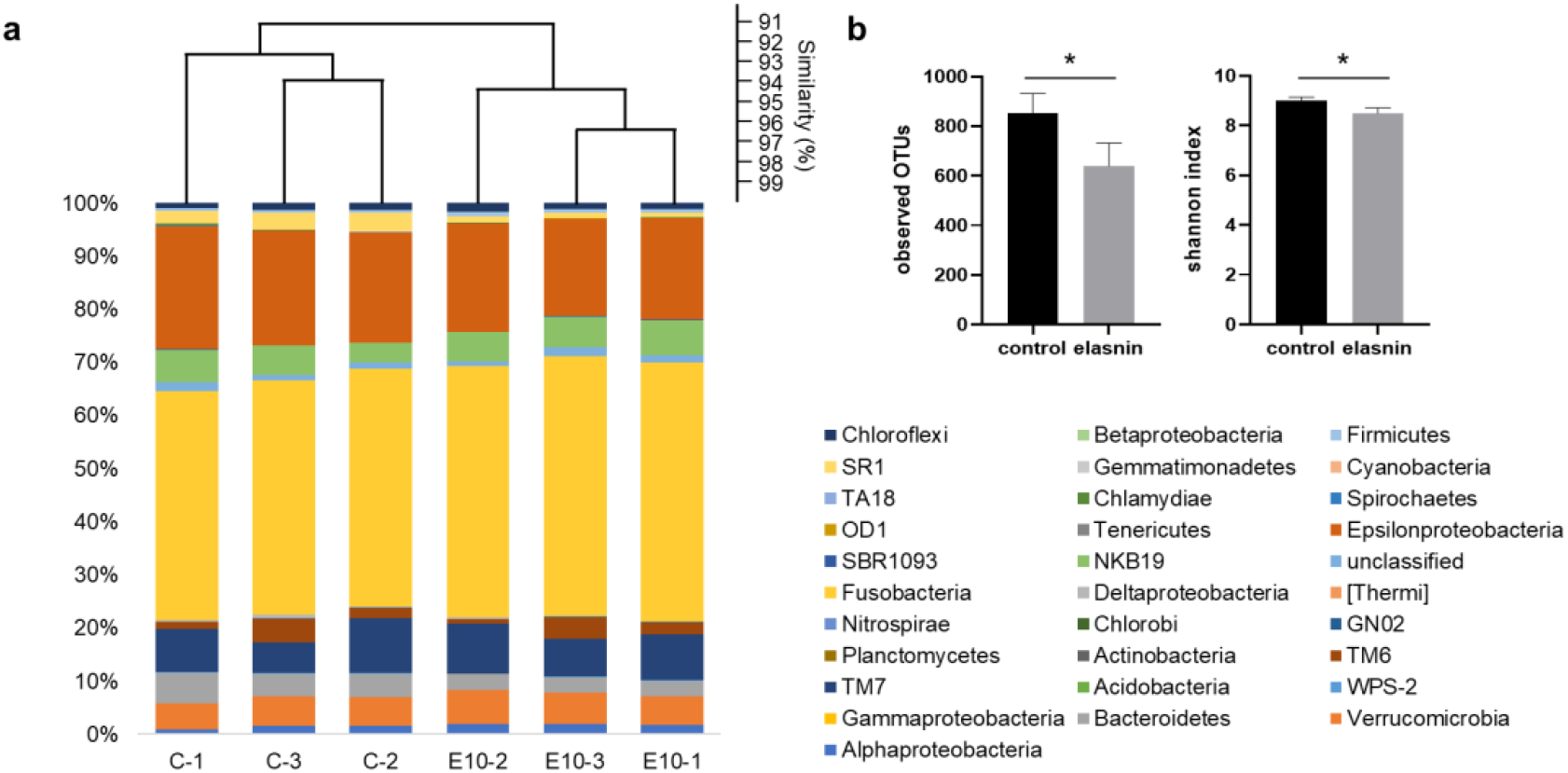
Composition analysis of biofilms grown on the control slides (without crude extracts) and treatment slides (with 10% CR coatings). **a** Similarity comparison of microbial compositions between biofilms on control slides (C-1,2,3) and 10 wt% elasnin-based coatings (E10-1,2,3) based on the beta-diversity (Bray-Curtis) at the phylum level. **b** Alpha-diversity of biofilms at the phylum level. The difference between the two types of biofilms was calculated by the student’s t-test and is indicated by an asterisk as follows: * for p<0.05.

## Discussion

The inevitable rise in microbial resistance and tolerance to antibiotics poses a serious threat to human health globally. Most of the persistent infections on human body surfaces and on medical devices are caused by biofilms^10,17^. Unfortunately, none of the antibiotics currently available in clinical treatment is designed to specifically target biofilm cells^18–21^. Herein, bioassay-guided isolation was applied to search for the compounds that specifically targeting biofilms. Among the four compounds with antimicrobial activity (Table 1) isolated in the present study, only elasnin exhibited antibiofilm activities. Comparing to the currently deployed antibiotic vancomycin, which failed to eradicate the mature biofilms, elasnin exhibited stable and high efficiency in inhibiting biofilm formation and eradicating mature biofilms of Gram-positive bacteria MRSA with low values of MBIC_90_ and MBEC_50_. Unlike vancomycin, which exclusively targets essential life processes and kills the pathogen, elasnin did not affect the cell viability and thus holds great potential in delaying resistance. Elasnin was first discovered in 1978 as a new elastase inhibitor with low toxicity in mice and high selectivity to human granulocyte elastase^22^. The low toxicity, biofilm-specific activity and selectivity of elasnin may make it a new alternative for treating biofilm-associated infections of Gram-positive bacteria, which prevents the development of antibiotic resistance.

**Table 1.** Bioassay results of the crude extracts from 12 actinobacterial strains

| Strain | Bioassay <sup>a</sup> |  |  | Identification |
| --- | --- | --- | --- | --- |
|  | MIC<br>(µg/mL) | MBIC<br>(µg/mL) | MBEC<br>(µg/mL) |  |
| *Streptomyces mobaraensis*<br>DSM 40847 | < 4 | < 4 | < 4 | Elasnin |
| *Nocardiopsis potens*<br>DSM 45234 | < 4 | > 100 | > 100 | Xanthone |
| *Streptomyces scabrisporus*<br>DSM 41855 | < 4 | > 100 | > 100 | Hitachimycin |
| *Streptomyces sulphureus*<br>DSM 40104 | 20-100 | > 100 | > 100 | Resistomycin |
| *Kutzneria albida*<br>DSM 43870 | 20-100 | > 100 | > 100 | - |
| *Streptomyces fulvissimus*<br>DSM 40593 | > 100 | > 100 | > 100 | - |
| *Streptomyces exfoliates*<br>DSM 41693 | > 100 | > 100 | > 100 | - |
| *Saccharothrix espanaensis*<br>DSM 44229 | > 100 | > 100 | > 100 | - |
| *Nocardiopsis synnemataformans*<br>DSM 44143 | > 100 | > 100 | > 100 | - |
| *Sciscionella marina*<br>DSM 45152 | > 100 | > 100 | > 100 | - |
| *Nonomuraea coxensis*<br>DSM 45129 | > 100 | > 100 | > 100 | - |
| *Streptomyces cattleya*<br>DSM 46488 | > 100 | > 100 | > 100 | - |

<sup>a</sup> All the crude extracts were tested with Gram-negative bacteria *E. coli* and Gram-positive bacteria MRSA; the concentrations illustrated here represent the bioassay results of the most potent crude extract among all.

The biofilm matrix is crucial to the resistance, persistence, and maturation of biofilms^2,27^, as the barrier formed by a large variety of EPS makes conventional antimicrobial treatment ineffective. Elasnin significantly inhibited cell aggregation and prevented the biofilm maturation. The potential of elasnin to destruct the biofilm matrix provides the basis for the development of new biofilm control strategies that could help fight to antibiotics resistance. Furthermore, elasnin destroyed the existed matrix, thus deactivated the protection of matrix to the cells and increased the cell exposure to environmental conditions, making them more susceptible to antibiotics. Combination therapy is considered as a new drug development strategy for combating antimicrobial resistance^23^ and elasnin’s interesting mode of action makes it an ideal candidate to be developed. It remains unknown how elasnin destroys the matrix. The investigation of elasnin’s mechanism at the molecular level may help to understand the biology of biofilms better and to identify some new antimicrobial targets.

Additionally, the wide type strain *Streptomyces mobaraensis* DSM 40847 produced elasnin in substantial quantity (0.33g/L) and was considered of great potential as a new industrial producing strain of antibiofilm agent. Low product yield has always been a limitation for the development and commercially applications of newly discovered biologically active compounds. Fermentation of this bacterium would solve the supply problem that often limits natural products from clinical and biotechnical development, paves the way for the later stage development of elasnin. Subsequently, elasnin-based antibiofilm coatings were easily prepared with low expenditures in the lab for the field test of antibiofilm efficiency to multispecies natural biofilms in the marine environment. The formation of marine biofilms on various submerged surfaces such as ship hulls and offshore infrastructures presents a serious problem, resulting in high economic loss every year. The selectivity of elasnin’s antibiofilm activity to Gram-positive bacteria also presents to uncertainty of elasin efficiency to multi-species biofilms. Surprisingly, elasnin-based antibiofilm coatings (10 wt%) remarkably inhibited the formation of multi-species natural biofilms that dominated by the Gram-negative bacteria (Fig. S4) for two weeks and exhibited antifouling activity in the fourth week. As revealed by 16S amplicon analysis, the introduction of elasnin caused a reduction in species richness and diversity of biofilms, which might be the reason for its antibiofilm and antifouling performances since signal molecules produced by the bacteria plays a key role in assisting the mixed biofilm formation and larval settlement. The changes in biofilm compositions may affect metabolic profile and therefore interfere with signals that mediate protist’s colonization, the settlement of invertebrate larvae and macroalgal spores^24–26^. The antibiofilm coatings began to lose their effectiveness after the third week in the field, probably due to the reduced release rate of elasnin. Different from those highly toxic antifouling paints that have been banned by the International Maritime Organization (IMO), elasnin is a natural product with low toxicity, therefore represents an ideal environmental-friendly antifoulant candidate ^27^. Collectively, the favorable antibiofilm and antifouling performance of elasnin-based antibiofilm coatings, combined with the low cost of supply and the low risk of environmental impact, could provide a new selection for the development of antibiofilm and antifouling materials.

In the present study, we identified a potent antibiofilm compound elasnin from the strain *S. mobaraensis* DSM 40847 in the course of our screening program of antibiofilm agents from twelve ‘genius’ strains of Actinobacteria Phylum. Elasnin exhibited biofilm-specific antibiofilm activities against Gram-positive bacteria with a low toxic effect and the ability of matrix destruction. With high productivity, elasnin-based antibiofilm coatings were easily prepared and presented a favorable performance in inhibiting the biofilm formation of natural multi-species biofilms and the attachment of macro-foulers in the marine environment. Overall, with low toxicity, low risks in resistance development and potential environmental impact as well as biofilm-specific activities, elasnin harbors great potential in the applications of biofilm control.

## Materials and methods

### Strains, culture media, and antibiotics

Twelve actinobacterial strains (Table 1) were purchased from the German Collection of Microorganisms and Cell Cultures (DSMZ, Braunschweig, Germany). The methicillin-resistant *S. aureus* ATCC 43300, *E. coli* ATCC 25922, and *S. aureus* ATCC 25923 were purchased from American Type Culture Collection. Marine *S. aureus* B04 was isolated from marine biofilms; both *S. aureus* B04 and *B. subtilis* 168 were obtained from the culture collection of our laboratory. Soybean powder was purchased from Wugumf, Shenzhen, China. Soluble starch was purchased from Affymetrix, Santa Clara, USA. Magnesium sulfate hydrate was purchased from Riedel-de-Haën, Seelze, Germany. Bacteriological peptone was obtained from Oxoid, Milan, Italy. Mueller-Hinton broth (MHB) was purchased from Fluka Chemie AG, Buchs, Switzerland. Phosphate-buffered saline (PBS) was purchased from Thermo Fisher Scientific Inc., San Jose, USA. Lysogeny broth (LB), glucose, 3-(4,5-dimethylthiazol-2-yl)-2,5-diphenyltetrazolium bromide (MTT), and 1-butanol were purchased from VWR International Ltd, Leicestershire, UK. Vancomycin and all other chemicals were supplied by Sigma-Aldrich Corporation, Saint Louis, USA.

### Compound isolation, productivity monitoring, and extraction efficiency comparison

Stock cultures of 12 *Actinobacteria* strains were inoculated into 50 mL of AM4, AM5 and AM6 media (Table S1) containing glass beads (to break up globular colonies) and incubated at 22°C and 30°C on a rotary shaker (170 rpm). The culture broth was extracted with 1-butanol on days 3, 5 and 7. The crude extracts were dissolved with DMSO before storage and bioassay. Pure compounds were isolated by reversed-phase high-performance liquid chromatography (HPLC) (Waters 2695, Milford, MA, USA) using a semi-prep C18 column (10×250mm) that was eluted with a 55-min gradient of 5–95% aqueous acetonitrile (ACN) containing 0.05% trifluoroacetic acid (TFA) at a flow rate of 3 mL/min. The structure of elasnin was elucidated through nuclear magnetic resonance (NMR) analysis of ^1^H, ^1^H-^1^H-COSY, ^1^H-^13^C-HSQC, and ^1^H-^13^C-HMBC NMR spectra recorded on a Bruker AV500 spectrometer (500 MHz) using dimethyl sulfoxide-d6 (^1^H-NMR DMSO-d_6_: δH = 2.50 ppm; DMSO-d_6_: δC = 39.50 ppm).

A stock culture of *S. mobaraensis* DSM 40847 was incubated in the AM4 medium as described in the section above. One mL of culture broth was taken every 12 hr and extracted with 1 mL of 1-butanol, ethyl acetate (EA) or n-hexane. The solvent for extraction was then removed by evaporation. The crude extract was dissolved in methanol and quantified through HPLC analysis with a Phenomenex Luna C18 column. The peak of elasnin was identified from the retention time, and its concentration was calculated based on an established standard curve.

### In vitro assays against single populations

MICs and MBCs were determined with methicillin-resistant *S. aureus* ATCC 43300 and *E. coli* ATCC 25922, according to the Clinical and Laboratory Standards Institute guideline CLSI M100 (2018). Briefly, a 10^5^ CFU/mL overnight culture of test strains was inoculated into MHB and treated with elasnin (or crude extracts and antibiotics) at a series of concentrations. After incubation for 24 hr, the minimum concentrations at which no bacterial growth was visible were recorded as the MICs. MBCs were measured following the MIC assay by plating 1 mL of suitably diluted culture broth from each well on Mueller-Hinton agar (MHA) plate. The MBC was defined as the lowest concentration at which an antimicrobial agent caused a more than 99.9% reduction in cells. Each assay was performed in duplicate and repeated three times.

MBICs and MBECs were determined as previously described ^28,29^. For the MBIC assay, an overnight culture of test strains was diluted into approximately 10^7^ CFU/mL with LB and 0.5% glucose and treated with various concentrations of elasnin (or vancomycin) in 96-well cell culture plates. Plates were then incubated at 37°C for 24 hr and rinsed twice with 1×PBS to remove non-adhering and planktonic cells. After rinsing, MTT staining assay was conducted to measure viable cells in the biofilms since MTT can react with activated succinate dehydrogenase in viable cell mitochondria to form blue-violet formazan, which can be read at 570 nm after dissolving in DMSO. For MBEC assay, an overnight culture of test strains was incubated for 24 hr in 96-well cell culture plates to form a biofilm but without adding elasnin. The formed biofilm was then rinsed twice with 1×PBS and challenged with elasnin at a series of concentrations (diluted with LB and 0.5% glucose) and incubated for another 24 hr at 37°C. After incubation, each well was rinsed twice with 1×PBS, and the MTT assay was conducted to measure viable cells in the remaining biofilm. Each assay was performed in quadruplicate and repeated three times; the lowest concentrations of elasnin that resulted in decreases of 90% and 50% in OD_570 nm_ were recorded as MBIC and MBEC respectively.

Methicillin-resistant *S. aureus* ATCC 43300 was used for the concentration-response curve study. A culture of 4×10^5^ CFU/mL MRSA in the exponential phase was inoculated into MHB with various concentrations of elasnin and vancomycin in 15 mL falcon tubes. Tubes were incubated at 37°C on a rotary shaker for 24 hr. One mL of culture broth in each tube was diluted with MHB, and 1 mL of diluted bacteria was plated on MHA plates for CFU counting. Culture broth from each well was inoculated on two plates, and the experiments were repeated three times.

### Coating preparation, field test, and release rate determination

A 4-L culture broth of *S. mobaraensis* DSM 40847 (incubated as described above) was extracted using n-hexane to obtain a sufficient amount of high-elasnin-content crude extracts. The elasnin-based antibiofilm coatings were prepared following the same procedures as those described in Ma et al. (2017). Typically, for the 10 wt% coatings, polymer (0.90 g, 90 wt%) and crude extracts (0.10 g, 10 wt%) were dissolved by vigorously stirring xylene and tetrahydrofuran (v:v=1:2) at 25°C. After enough mixing, a glass slide was coated with the solution and left to dry at room temperature for a week to remove the solvent. The same procedure was followed to prepare coatings with different concentrations of crude extracts.

Coated glass slides were submerged in seawater at a fish farm in Yung Shue O, Hong Kong (114°21′E, 22°24′N) for two to four weeks. Afterward, the glass slides were retrieved and transported back to the laboratory in a cooler with *in situ* seawater and were washed twice using autoclaved and 0.22 μm filtered seawater to remove loosely attached particles and cells. The slides were then stained using the FilmTracer™ LIVE⁄DEAD Biofilm Viability kit and examined under confocal laser scanning microscopy. At the same time, the release rate of elasnin was determined by measuring its concentration using HPLC under static conditions. The coated panels were immersed in 100 mL of sterilized artificial seawater held in a measuring container. Ten mL of seawater was taken after immersion for 24 hr and elasnin was extracted with the same volume of dichloromethane, which was then removed under nitrogen gas. After drying, the extract was resuspended in 100 mL of methanol and underwent HPLC analysis as described above. The release rate was measured every week for four straight weeks and each concentration was tested in duplicate.

### DNA extraction, 16S rRNA gene sequencing, and analyses

Biofilm samples on the coated slide surface were collected with autoclaved cotton and stored in DNA storage buffer (10 mM Tris-HCl; 0.5 mM EDTA, pH 8.0) at −80 °C. Before the extraction, samples were vortexed several times to release the microbial cells into the DNA storage buffer. All the samples were then subjected to centrifugation at 10,000 rpm for 1 min, and the supernatant was discarded. After continuous treatment with 10 mg/mL lysozyme and 20 mg/mL proteinase K, DNA was extracted from the treated microbial cells with a microbial genomic DNA extraction kit (Tiangen Biotech, Beijing, China) following the manufacturer’s protocol.

The quality of DNA samples was controlled using NanoDrop (which tests the DNA purity, OD260/OD280) and agarose gel electrophoresis (which tests for DNA degradation and potential contamination). The hypervariable V3-V4 region (forward primer: 5 ′ -CCTAYGGGRBGCASCAG-3 ′ ; reverse primer: 5 ′ - GGACTACNNGGGTATCTAAT-3′) of prokaryotic 16S rRNA genes was used to amplify DNA from biofilms by polymerase chain reaction (PCR). PCR products were purified before library construction and sequenced at Novogene (Beijing, China) on the NovaSeq 6000 System. The read length was 250 bp, and each pair of reads had a 50-bp overlapping region. The paired-end reads were subjected to quality control using the NGS QC Toolkit (version 2.0). The 16S rRNA gene amplicon data were analyzed using the software package QIIME2 then merged using Q2_manifest_maker.py in QIIME2. The low-quality reads and chimeras were removed using dada2 commands in QIIME2. To normalize the uneven sequencing depth, 500,000 filtered reads for each sample were picked up. Operational taxonomic units (OTUs) were classified de novo from the pooled reads at 97% sequence similarity using a classifier trained by the Naive Bayes method. Representative sequences were then recovered using the feature-classifier classify-sklearn script in QIIME2. The alpha-diversity analyses (observed OTUs and Shannon diversity) were performed using the script ‘qiime diversity alpha’ in QIIME2. Beta-diversity based on the Bray-Curtis distances was conducted by the cluster analysis in the software PAST (version 3.0). Additionally, the taxonomic structure was drawn in Excel (Office 365 MSO 64-bit) based on the relative abundance.

### CLSM observation with biofilm cell and matrix staining

Biofilms were grown on glass cover slides as those described for the MBIC and MBEC assays. Treated biofilms were then rinsed twice with 1 × PBS and stained with FilmTracer™ FM® 1-43 green biofilm cell stain and FilmTracer™ SYPRO® Ruby Biofilm Matrix Stain at room temperature for 30 min in the dark. Leica Sp8 Confocal Microscope was employed to observe cells and matrix in the biofilm at 488 nm.

### Statistical analyses

Statistical analyses for all data were performed using the GraphPad Prism 8.0.2 software. The composition of the biofilm on the coatings was compared with that in control groups using the student’s t-test.

## Acknowledgment

This work was financially supported by the National Key R&D Program of China (2018YFA0903200), China Ocean Mineral Resources Research and Development Association (COMRRDA17SC01) and the Hong Kong Branch of Southern Marine Science and Engineering Guangdong Laboratory (Guangzhou) (SMSEGL20SC01).

